# High-Definition Electronic Genome Maps from Single Molecule Data

**DOI:** 10.1101/139840

**Authors:** John S. Oliver, Anthony Catalano, Jennifer R. Davis, Boris S. Grinberg, Timothy E. Hutchins, Michael D. Kaiser, Steve Nurnberg, Jay M. Sage, Leah Seward, Gregory Simelgor, Nathan K. Weiner, Barrett Bready

## Abstract

With the advent of routine short-read genome sequencing has come a growing recognition of the importance of long-range, structural information in applications ranging from sequence assembly to the detection of structural variation. Here we describe the Nabsys solid-state detector capable of detecting tags on single molecules of DNA 100s of kilobases in length as they translocate through the detector at a velocity greater than 1 megabase pair per second. Sequence-specific tags are detected with a high signal-to-noise ratio. The physical distance between tags is determined after accounting for viscous drag-induced intramolecular velocity fluctuations. The accurate measurement of the physical distance between tags on each molecule and the ability of the detector to resolve distances between tags of less than 300 base-pairs enables the construction of high-density genome maps.

## Introduction

In the last decade, the development of novel methods of DNA sequencing has dramatically lowered per base costs while providing enormous gains in throughput. Most platforms have short read lengths, limiting the overall usefulness of the data, particularly when dealing with highly repetitive genomic sequences. While small repeats can be spanned directly by emerging long-read technologies or synthetically by paired-end approaches, large or highly complex repeated sequences remain a significant challenge during assembly.^1^ In addition, current sequencing and assembly methods remain unable to accurately and consistently resolve structural variants.^2^ This is of particular concern for the clinical application of structural variant detection.

Physical mapping provides information over a larger scale and has been used for decades to aid in the assembly of sequence information. The generation of genetic and physical maps was an important milestone in the assembly of a human reference genome by the public consortium.^3^ More recently physical maps have been used to aid in assembly projects with genomes ranging from bacteria to human.^4,5,6^

Existing mapping technologies rely on optical imaging of long DNA molecules with sequence specific fluorescent labels. While providing excellent long-range information, the nature of optical detection imposes physical limits on the accuracy and resolution of the data. The relatively low sensitivity of optical detection methods also impedes data acquisition speeds and thus the throughput of detectors.^7^ The use of an electronic detection scheme in contrast has several advantages over optical mapping. This system does not suffer from optical resolution limits and allows for increased sensitivity with lower cost, maintenance, and space requirements in comparison to optical systems.

For over two decades, nanopores have been used as highly sensitive single-molecule electronic detectors.^8,9,10,11^ Both solid-state and protein pores have been envisioned as detectors to enable rapid DNA sequencing.^12^ Most of the proposed nanopore-based sequencing methods rely on the sequential identification of individual bases in a DNA strand as they translocate through the pore^13,14,15,16^ or by the detection of base-specific tags that are released during sequencing by synthesis occurring near the pore entrance.^17^ Most systems rely on an ionic current blockade by the analyte for detection. The degree to which the flow of ions through the nanopore is changed is dependent on the size of the molecule, how effectively it interferes with the movement of ions through the detector, and on the occupancy time of the molecule in the detector. While detection of groups of nucleotides in a DNA strand has been accomplished and the sequence can subsequently be deconvoluted, the reliable determination of single base-resolved sequence in the nanopore comes with a reduced velocity of translocation.^18^

Solid-state nanochannels have a number of advantages over nanopores while retaining the high sensitivity afforded by electronic detection when appropriately designed. Among the advantages are the ability to control electric field gradients in front of the detector, the ability to control the conformations of molecules in the device, the ability to multiplex detectors and control crosstalk without electrically isolating neighboring detectors, and the ability to easily fabricate large numbers of devices in semiconductor foundries.

Here we describe a novel physical mapping platform, Nabsys *HD-Mapping™*, which utilizes a nanochannel configuration to electronically detect single long DNA molecules that are tagged in a sequence selective fashion. The tagged molecules are translocated through the nanochannel detector and the relative positions of the subsequences are determined. The data characteristics of this approach will be presented with an emphasis on precision, positional accuracy, and resolution of sequence-specific tag detection.

A photograph of the electronic mapping instrument is shown in Figure 1. The housing has 4 ports on top that accommodate standard laboratory pipet tips for introduction of buffers and samples. A fluid manifold connects the pipet ports to the detector and contains driving electrodes that are used to electrophorese samples through the detection region. Detector chips are inserted into the housing and a hinged lid seals the detector to the fluid manifold. The instrument contains all of the requisite electronics for signal amplification and output to a computer.

**Figure 1.**
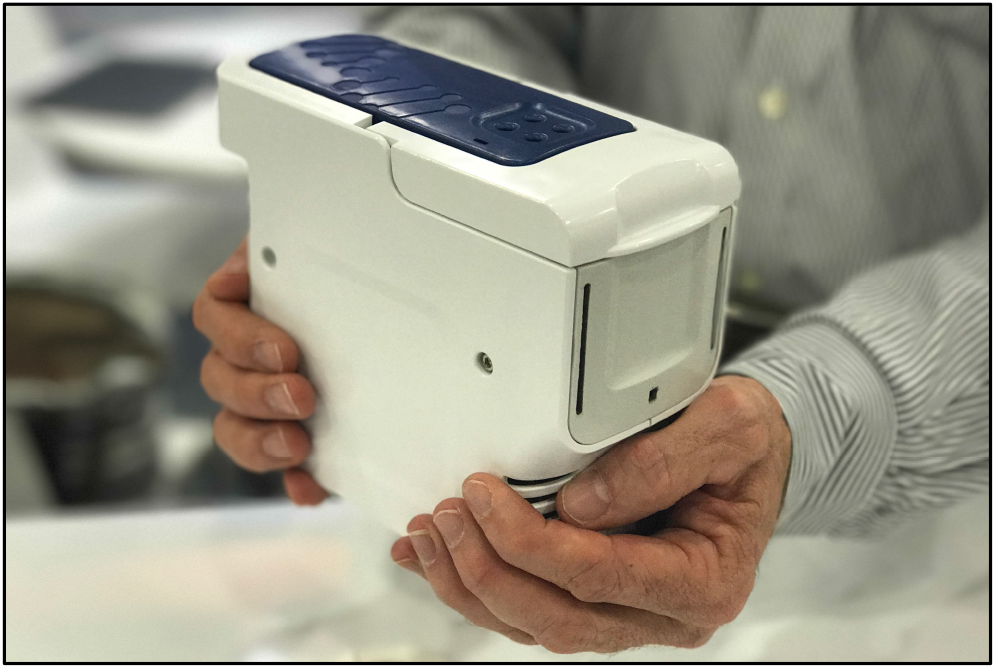
The Nabsys electronic mapping instrument. The instrument shown contains a detector, fluidic manifold that allows standard laboratory pipets to introduce sample or buffers, pumps to introduce samples and buffer to the detector chip, and all electronics required for signal amplification. The control software and signal processing software reside on a laptop or desktop computer.

### Detectors

The nanochannel detector described here overcomes the limitations of nanopore detection. By defining the geometry of micro- and nanochannels around the detector, the conformation of DNA strands entering the detector is controlled. The geometry is also used to control the electric field and maintain conditions of free solution electrophoresis to bring molecules to the detector. Under conditions of free draining electrophoresis, the rate of transport of DNA is independent of the length of the polymer. As a result there is no length selection prior to detection and the distribution of DNA lengths observed by the detector is representative of the sample size distribution.

The entrance geometry of the nanochannel detector is designed to create field gradients that elongate the DNA prior to entering the detector. Individual molecules have similar conformations during the approach to the detector and thus a smaller variation in intermolecular velocities than has been observed for nanopores.^19^

The access resistance and the resistance between multiple detectors accessed by one microchannel are controlled by the microchannel cross-sectional areas. The perturbation of the nanochannel resistance during DNA translocation is a small percentage of the total resistance. The high inter-detector resistance combined with the small resistance change of the detector during event detection provides a system that can be multiplexed from the same driving electrodes and microchannel without crosstalk.

Nanopores and nanochannels, while both enabling electronic detection, differ in several key factors. One downfall of the nanopore geometry for the direct analysis of biological polymers is that prior to threading into the nanopore, there is no control of the DNA conformation. As a result, there is a significant barrier to capturing the end of a DNA strand.^20^

The translocation behavior after threading is also complicated by the variety of conformations accessible to the polymer.^21^ In addition, both solid-state and biological nanopore DNA detection schemes typically utilize a geometry in which the pore diameter is significantly occluded during detection of the DNA analyte. While this geometry maximizes the observed signal, it results in significant changes in the resistance of the system. If multiple pores are placed in parallel and they are not electrically isolated from one another, the current blockade occurring in one pore significantly affects the current flowing in the other pores. In order to multiplex nanopores, they must be electrically isolated.

A schematic of the Nabsys detector is shown in Figure 2. The detector comprises ports for sample and buffer introduction, microfluidic channels for delivery of sample to the detection region, a nanochannel detector, and electrodes for sensing the potential drop in the nanochannel. Electrodes fluidically connected to the ports are placed outside the device to provide an electrophoretic driving force and serve as the source for the potential drop in the nanochannel. A fluid manifold is routinely connected to the ports to allow sample and buffer introduction with standard laboratory pipets.

**Figure 2.**
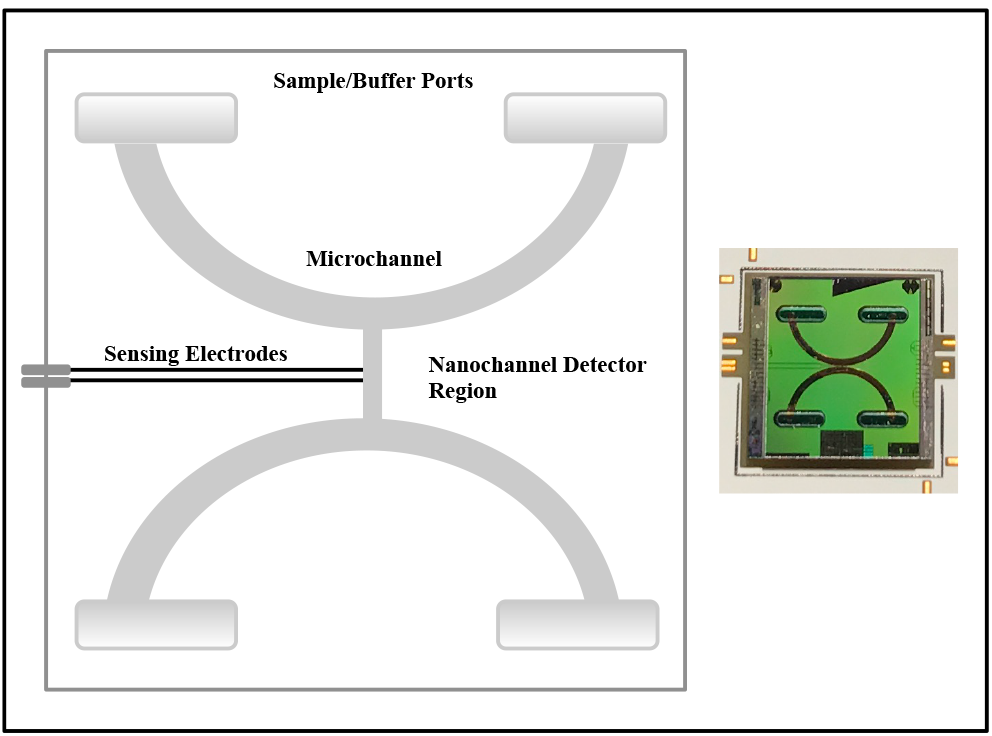
Detector design. A schematic of the detector showing ports for introduction of buffer or sample, microchannels for delivery of analytes to the detector, the detector and associated sensing electrodes.

### Sample Preparation

Relative to other mapping approaches, Nabsys’ electronic detection method is able to resolve closely spaced tags, down to a few hundred base pairs apart, and can accurately measure distances between tags, resulting in higher average information density. One important effect of this increased information content is the ability to utilize DNA lengths ranging from 30 kilobase pairs (kbp) to 100’s of kbp depending on the genome size. As a result, solution phase kit-based DNA extraction procedures can yield DNA of sufficient length for assembly and structural variant detection applications. In the Nabsys platform, following DNA isolation, samples are nicked with one or a combination of commercially available nicking endonucleases. Nicking is followed by tag attachment prior to coating with *E. coli* RecA protein, a bacterial DNA binding protein that uniformly coats double-stranded DNA (dsDNA).

While the translocation of naked DNA through nanopores has been widely studied,^22^ the translocation of RecA-coated DNA has only recently appeared.^23,24,25^ RecA coating prior to translocation provides several benefits for single-molecule analysis of DNA. dsDNA has an average diameter of 2.2 nm while the diameter of RecA-coated dsDNA has been estimated to be 8-10 nm.^26^ The larger cross-sectional area of the nucleoprotein filaments increases the signal-to-noise ratio during detection. RecA coating also increases the persistence length of the DNA. Naked dsDNA behaves as a polymeric chain with a persistence length of approximately 50 nm in typical buffer solutions while RecA-dsDNA nucleoprotein filaments have a persistence length of 962 ± 57 nm.^27^ This increase in persistence length relaxes the requirements on the device geometry used to linearize molecules upstream of the detectors.

As single molecules translocate through the nanochannel detector, the resistance increases for the duration of the translocation. The magnitude of the change in resistance is proportional to the volume of the detector that is occupied by the analyte analogous to blockade detection in nanopores, which has been well described.^28^ When a molecule enters the detector, the baseline potential between the sensing electrodes rises to a new stable level and stays at the new level until the molecule exits the detector at which time it returns to the baseline level. The time of the start and end of each event is recorded as well as the magnitude of the change in potential of the detector.

When DNA is tagged, single-molecule events are detected as they transit through the detector. A resistance change of the detector that is equal in magnitude to the blockade observed with untagged DNA indicates the entry of a single molecule into the detector. In contrast to untagged events, a tagged event is characterized by additional short duration blockades residing on top of the continuous blockade created by the DNA molecule. These short blockades denote the translocation of a tag that is attached to the DNA, through the detector. An example of a single DNA molecule that has been tagged is shown in Figure 3.

**Figure 3.**
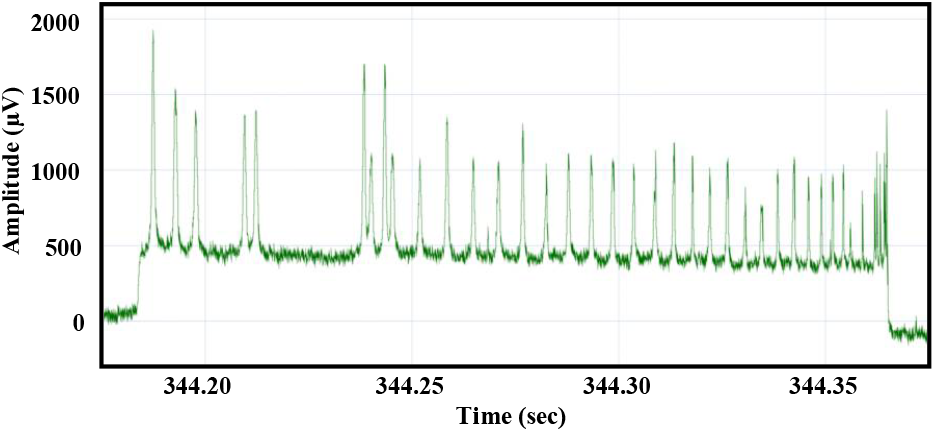
A representative single molecule event collected during translocation of tagged DNA. The DNA molecule shown is 165 kbp in length.

### Molecule Velocity

In contrast to imaging static DNA molecules, the translocation of DNA through nanopores or nanochannels is characterized by inter- and intramolecular velocity changes that must be accounted for during processing of the raw signal. Both molecular conformation and drag have been suggested as a cause for the observed variations in velocity.^21,29,30^

In the Nabsys system, the initial conformation of the molecule has very little effect on the variation of velocity. The high persistence length of the nucleoprotein molecules and the high degree of confinement during the approach to the nanochannel detector stretch molecules into a linear conformation prior to entry. We have previously described the effect of viscous drag on intramolecule velocity during translocation through solid-state nanopores.^31^ The same viscous drag model also describes the intramolecular change in velocity that we observe in our nanochannel detectors.

In the context of the Nabsys mapping platform, determining the instantaneous velocity of DNA molecules in the detector is essential for converting the interval time measurements to DNA molecule lengths. The time-to-distance transformation must be valid for all lengths of DNA since the platform uses long DNA from random fragmentation of genomes. As we have previously described for nanopores, the dominant forces acting on DNA molecules as they translocate through our nanodetector are provided by i) the constant electric field inside the detector and ii) the variable viscous drag resulting from the interaction of the fluid with the portion of the molecule that has not yet entered the detector. The velocity of molecules in the detector results in a very low Reynolds number which implies purely laminar flow. A Stokes’ drag model is therefore appropriate to model the behavior of translocating molecules. When the velocity profile analysis is applied to events representing the translocation of molecules with bound tags, the physical distance between the tags on each molecule is determined accurately. These transformed events may then be used to assemble maps or align events to a known map.

The Nabsys platform has a number of advantages for the determination of both inter- and intra-molecule velocity changes. First, the length of DNA that is routinely used in our system is much longer than that used in typical nanopore studies allowing a wider variation in intramolecular velocity to be observed. Second, our system has a narrow distribution in translocation times for molecules of equivalent length so that small changes in intramolecule velocity can be accurately characterized. Third, our sample preparation method allows high-density DNA tagging, enabling velocity measurements on many intervening segments between tags on a given molecule. The variation in instantaneous velocity during the translocation of each molecule can be observed when molecules with tags at known positions are translocated and the time between the tags is analyzed.

Analysis of the tagged translocation events confirms that the model we have previously described for translocation of RecA-coated DNA through solid-state nanopores holds for the detectors described herein. In brief, the observed behavior is a result of the effect of Stokes’ viscous drag acting on the portion of the molecule that has yet to be translocated through the detector. A model that considers two forces acting on each molecule, an electrophoretic force in the direction of translocation that is constant and a drag induced force acting in the opposite direction that changes with the length of the molecule that has not yet passed through the detector, accounts for the observed velocity profiles.

The general form of the relationship of translocation time and the length of the molecule is:

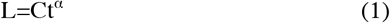

where L is the length of the molecule in base-pairs, C is a constant, t is the time of translocation and alpha is an exponent that captures the conversion of the volume of the nucleoprotein filament to an equivalent volume sphere. Experimentally, with DNA lengths ranging from 10 kbp to greater than 200 kbp we find that alpha is 0.56 in our system.

## Results

### Detector Characterization

Phage lambda DNA (48.5 kbp) provides an illustrative example of the effect of viscous drag on molecules as they translocate through a Nabsys nanochannel detector. There are ten recognition sites for nicking endonuclease Nt.BspQI in the phage lambda genome. The relative positions of the nicking endonuclease recognition sites are indicated by vertical lines in the schematic representation in Figure 4. When lambda DNA is nicked and then tagged with proprietary chemistry, each full-length molecule is expected to have ten tags attached. The size of each interval, defined as the distance between consecutive tags, is indicated in base pairs in Figure 4.

**Figure 4.**
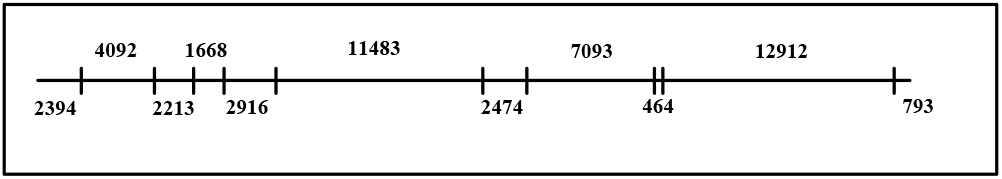
Representation of the ten Nt.BspQI nick sites on phage lambda. The distance between each tag and to the beginning and the end of the molecule are indicated in base pairs.

Displaying translocation times of events in the format of a histogram is useful in evaluating sample length distribution as well as translocation time variation in a single population. A histogram of the translocation time for all events longer than 5 ms from a representative run of tagged lambda is shown in Figure 5. A major peak centered at 36 ms is present in the histogram. This peak primarily represents full-length tagged lambda DNA. Shorter duration events were also observed. The shorter events were presumed to be primarily fragmented lambda DNA.

**Figure 5.**
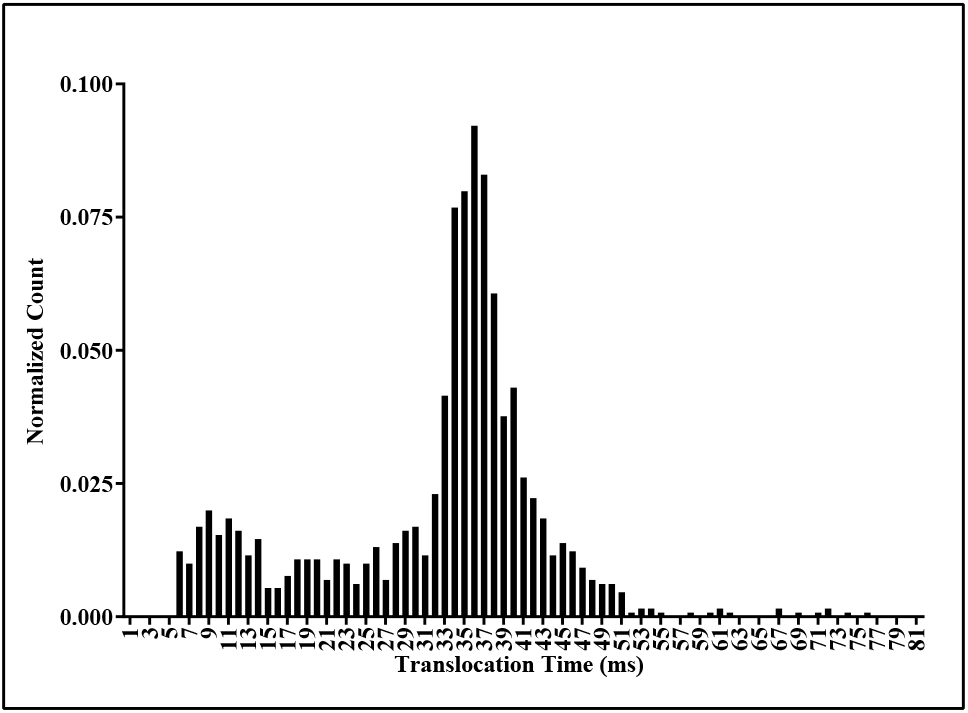
Histogram of the frequency of observed translocation times for phage lambda DNA tagged at Nt.BspQI sites (n=1302).

The data from the same run of tagged lambda were filtered by selecting only those events containing 9, 10, or 11 tags. The filtered events include full-length molecules that may have a single false positive or false negative as well as shorter molecules that happen to include the requisite number of tags. A histogram of translocation times of the filtered data is shown in Figure 6. A dominant peak corresponding to tagged full-length phage lambda is observed in the chart. The mean translocation time for the collection of all events is 36.4 ± 6.5 ms and the full width at half maximum of the major peak is 6 ms. The tight distribution of translocation times is a reflection of the uniform behavior of molecules in the detector. The Nabsys platform also displayed a high level of precision between different detectors. The mode of the large peak had an average value of 37.6 ± 2 ms for 10 different detectors (data not shown).

**Figure 6.**
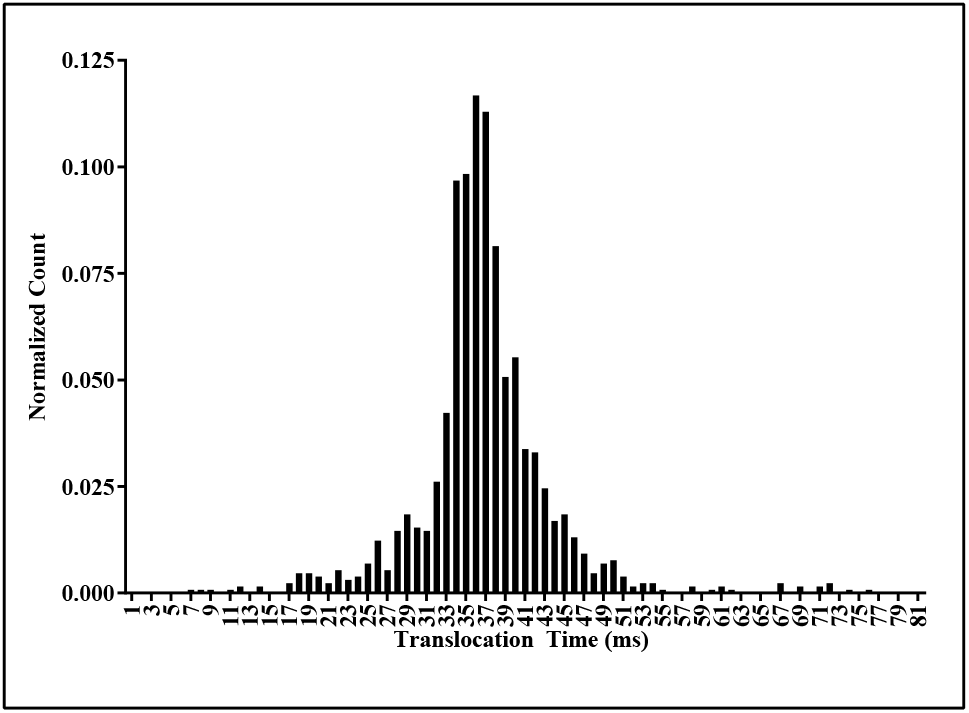
Histogram of the frequency of translocation times observed for phage lambda DNA containing 9-11 tags (n=773).

The intramolecular variation in velocity can be quantified by considering the distribution of times between tags on full-length molecules. An examination of the data showed two apparent populations based on the observed pattern of tags.

Lambda phage DNA is characterized by an asymmetric pattern of recognition sequences for Nt.BspQI (Figure 4). Therefore, upon detection, tagged lambda molecules display two different patterns of tags, dependent on entry into the detector in either of the two orientations. Both populations have the same distribution of total translocation time. Thus the average velocity was the same for each molecule and was independent of the entry direction of the molecules into the detector.

Representative examples of each of the two populations are shown in Figure 7. Events in each group had the same profile of resistance versus time demonstrating that the translocation of molecules was consistent within each population. If the intramolecular velocity were invariant during translocation, the events in one population would be expected to be mirror images of the events in the other population. Figures 7a and 7b are representative events from each of the two populations. Figure 7c is a mirror image of the event in Figure 7a.

**Figure 7.**
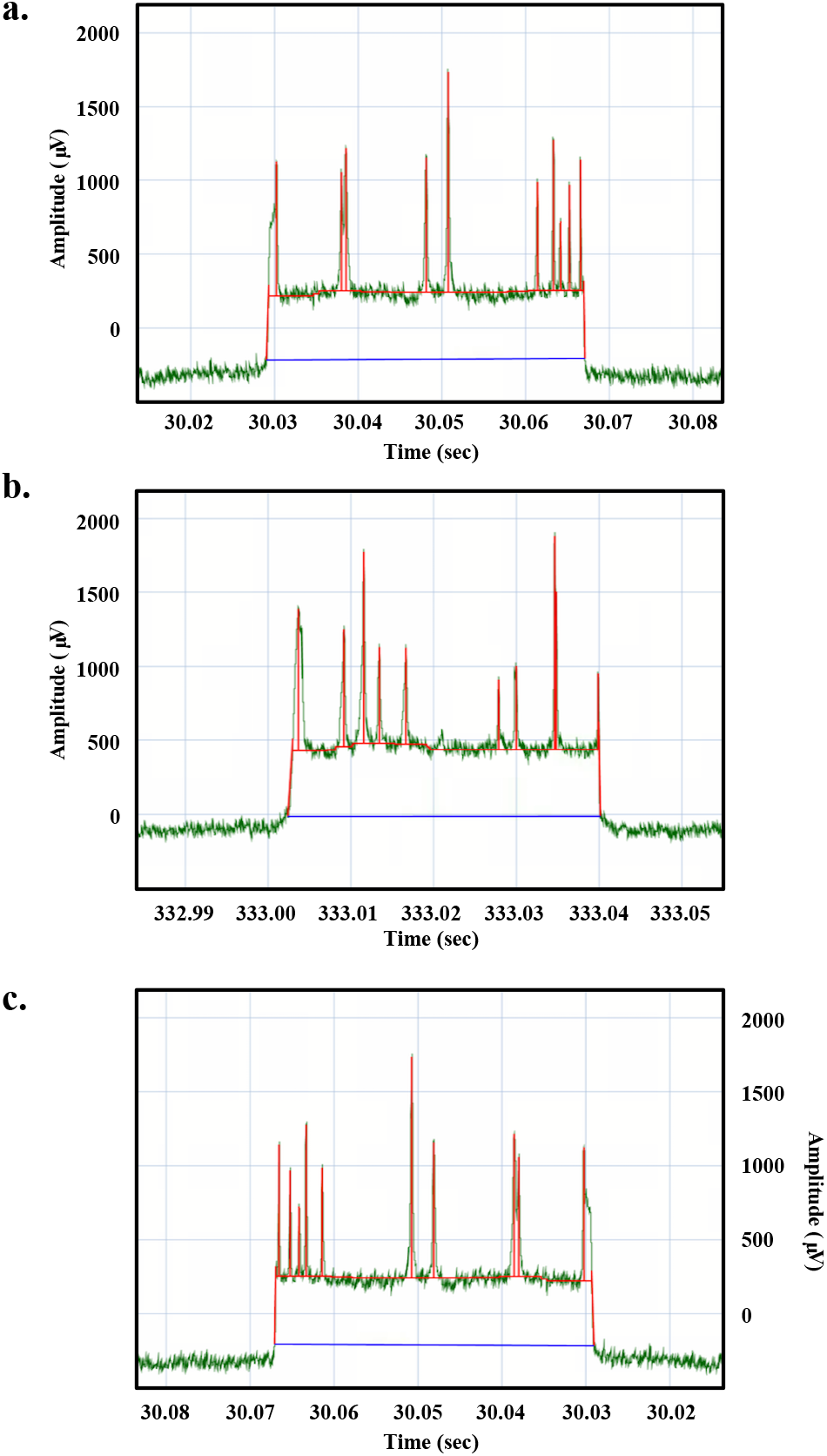
Representative examples of tagged lambda DNA translocation blockade events from each of two populations. The raw event is shown in green. Signal processing interpolates a baseline under each event shown in blue and picks events and tags shown in red. The signal arising from translocation of a representative event through the detector in (a) population A, and (b) population B are shown. The mirror image of the event in (a) is shown in (c).

According to the prediction stated previously, if the velocity of the molecules was constant during translocation, the events in Figures 7b and 7c would look similar. Inspection of the images in Figures 7b and 7c reveals that the two images are not superimposable based on the time of appearance of corresponding tags in each molecule. Thus, the velocity of the molecules must change during translocation.

Applying the Stokes’ drag model previously described to each event starting from the end of the event corrected for the intramolecular velocity change and was used to transform time dependent measurements into distance. If the detector was calibrated with a DNA sample of a known single length (in this case full-length lambda DNA is assumed to have a length of 48.5 kbp) the value of the constant C for expression of length in base pairs in Equation (1) could be determined.

Transformation of the data from the time domain to distance corrected for velocity changes and converted all time intervals in the molecules to a distance measured in base pairs. Single-molecule events representing full-length lambda DNA still had two populations of tag patterns. However, the populations were now mirror images of one another. This indicated that the existence of two distinct populations was due to the direction of entry into the detector and that the transformation had corrected for the intramolecular velocity changes. A histogram of the transformed phage lambda data is shown in Figure 8. The mean length of the events is 48.5 ± 4.7 kbp.

**Figure 8.**
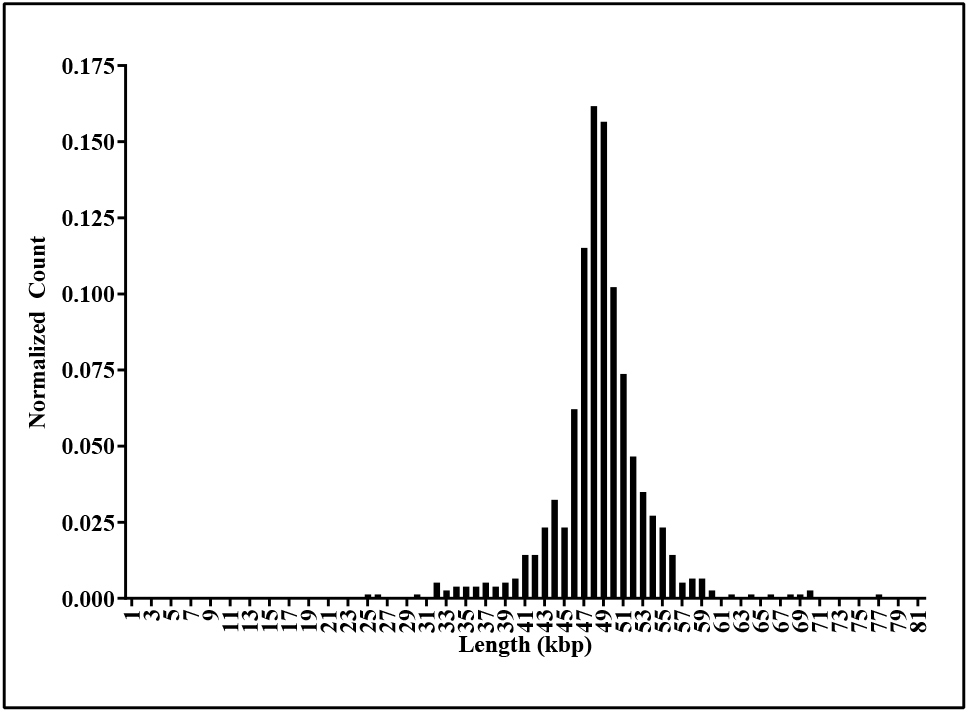
Histogram of the frequency of lengths observed for mapped phage lambda DNA molecules containing 9-11 tags (n=773).

Representations of a portion of the single-molecule temporal events, transformed to distance, are shown in Figure 9. Events colored in black entered the detector from the left side of the molecule as displayed (population A); blue events entered the detector from the right side of the molecule as displayed (population B).

**Figure 9.**
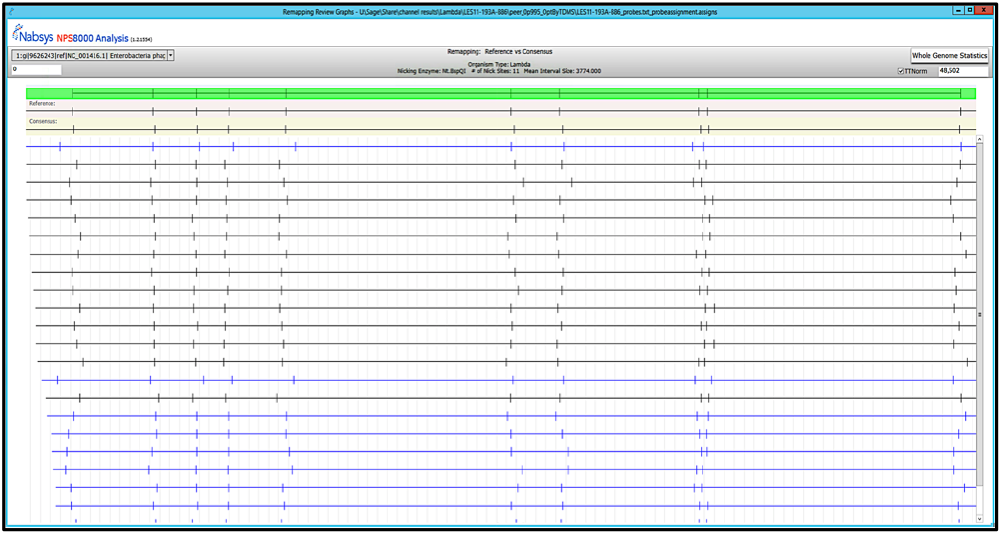
Tag locations in single molecules detected by the Nabsys platform for Nt.BspQI nicked phage lambda DNA. Molecules that enter the detector from either end have the same pattern of probes after transformation from the time domain to the spatial domain with Equation 1.

The data was further filtered by selecting only those events that have tags at each end of the event. The position of each tag on every molecule was determined. The mean of the measured tag positions for the molecules, grouped by translocation orientation, is shown in Table 1. The difference in calculated positions for each tag in the two populations is shown in the last column. There is close agreement between the calculated positions of the tags for each of the two populations implying that no directional bias is present. Further confirmation depends on the analysis of more complex samples that are composed of variable length molecules having more varied and complex tag patterns.

**Table 1.**
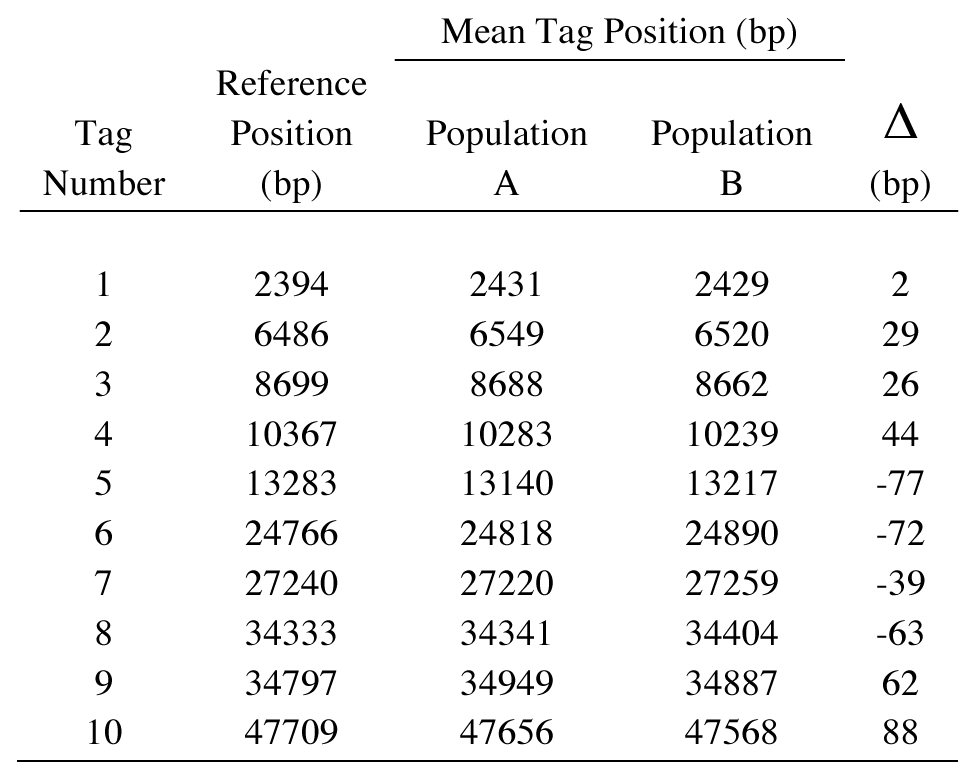
Mean tag positions for each of the ten sites in Nt.BspQI nicked phage lambda DNA (n=703).

### Mapping of a Bacterial Genome

Accurate molecule velocity correction also enabled more complex, randomly fragmented samples to be analyzed. As an example, data were collected from a bacterial genome. The data were subsequently both mapped to a reference map and *de novo* assembled into a genome map. The quality of the data and of the resulting assembly was determined by comparing to a reference sequence.

Genomic DNA was isolated from *Escherichia coli* (*E. coli*) MG1655 using a solution-phase DNA isolation kit. The genomic DNA was nicked with endonucleases Nt.BspQI and Nt.Bpu10I in separate reactions with average interval sizes predicted to be 6.8 kbp and 3.9 kbp respectively for the two reactions. The interval size distributions for nicking with the two enzymes and the percentage of those intervals that are < 300 bp, 300-1500 bp, and > 1500 bp are shown in Figure 10. In contrast to optical based mapping approaches where the resolution is 1500 bp – 2000 bp,^32,33,34,35,36,37^ the Nabsys detector was able to resolve intervals below 300 bp. The resolution of the detector enables the use of a wide variety of nicking enzyme densities.

**Figure 10.**
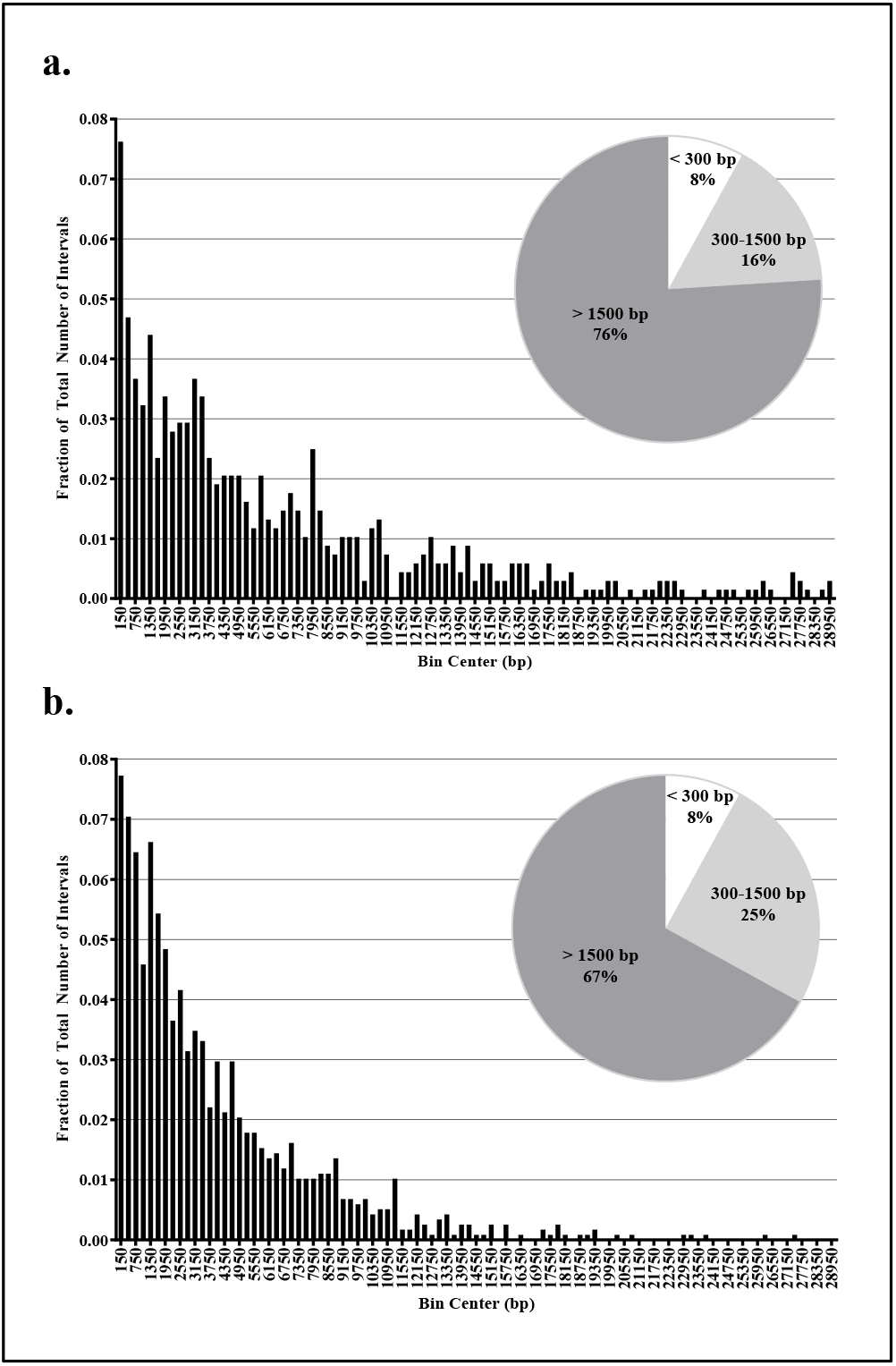
Reference interval size distribution histograms. (a) Nt.BspQI and (b) Nt.Bpu10I nicking of *E. coli* MG1655. Inset graphs indicate the percentage of intervals that are < 300 bp, 300-1500 bp, and > 1500 bp.

The tagged samples were separately introduced into detectors. The platform collects data unattended. During data collection, statistics on the event frequency, event length distribution, and baseline stability are recorded. If the detector clogs as determined by either low event frequency or a noisy baseline, the platform automatically performs a declog procedure. Declogging of the detector does not perturb the sample. Once the detector has been declogged, the run continues. Signal processing is performed during the run, and the run can be controlled and observed either from an attached computer or from devices that can remotely connect such as a laptop or a smartphone.

Each single molecule event obtained during data collection was transformed from the temporal domain to the spatial domain. An event may contain one or more source(s) of error, including: interval size error, false positive tags (tags present on the event, but absent in reference) and false negative tags (tags present in reference, but absent on the event).

False negatives may arise from nicking errors, incomplete tagging, signal processing errors, and alignment errors. False positives may arise from offsite nicking, co-translocation of molecules, signal processing errors, or alignment errors. We found that both false negatives and false positives were stochastic and are rapidly eliminated from consensus maps at very low coverage. The stochastic nature of the false positives eliminates consistent off-site nicking by the nicking enzyme, or star activity, as a potential source of false positives.

For each nicking endonuclease, a reference map was created by locating the nicking endonuclease recognition sequence, or its reverse-complement,^38^ within a reference FASTA file. The resulting reference map contained the expected tag locations on each chromosome.

A proprietary seeding algorithm was used to determine all feasible alignments between each read and the reference. Seeding stringency was set to eliminate infeasible alignments while preserving >95% of all correct alignments.^39^ With large reference maps, the use of seeding improves remapping speed by several orders of magnitude. A proprietary scoring function, in conjunction with dynamic programming, was used to determine the highest likelihood alignment between each read and the reference. The resulting score for each read was evaluated with a proprietary thresholding function to determine whether the read adequately matched the reference and had sufficient information content to be included in the set of remapped reads. The set of remapped reads was then used to construct a consensus map.

Single-molecule electronic mapping data were collected with each nicking approach and mapped to their respective reference maps to an average coverage depth of ~250x. The single-molecule false negative rate was 3.3% and 2.8% for Nt.BspQI and Nt.Bpu10I respectively. The single-molecule false positive rate was ~1 in 300,000 bp for both samples.

Spanned interval coverage maps of the two data sets are shown in Figure 11. The data have a relatively even depth of coverage across the 4.6 Mbp genome with normalized coverage standard deviations of 0.24 and 0.43 fold coverage respectively for Nt.BspQI and Nt.Bpu10I.

**Figure 11.**
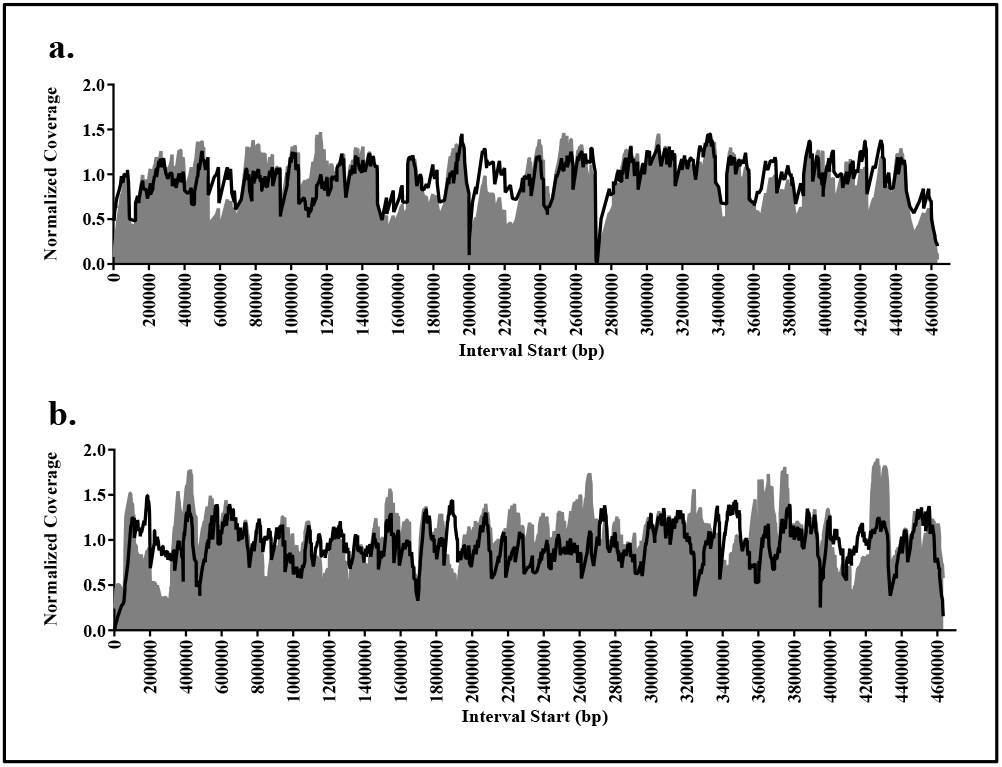
Spanned interval coverage maps for (a) Nt.BspQI and (b) Nt.Bpu10I normalized for average coverage of each data set. Overlaid black lines indicate the predicted coverage variation due to effects of nick density variation and potential proximity breaks.

The moderate variation in coverage observed locally is largely due to variation in regional nick site density throughout the genome. In addition, melting of the DNA strands between nick sites that are in close proximity to each other on opposite strands can lead to a DNA break. The impacts of local nicking density and coverage loss due to proximity breaks are well-understood phenomena allowing for the accurate prediction of local coverage depth for a given data set with a known reference map as shown by the black line in Figure 11.

For each of the two samples, the reference interval size vs. the consensus interval size was plotted. The plots for each nicking enzyme are shown in Figure 12. In both cases the consensus interval sizes are in good agreement with the reference interval sizes as shown by the quality of the fit. This relationship extends down to ~300 bp, well below the optical diffraction limit (Figure 12 (a, b)) highlighting the enhanced resolution inherent in the Nabsys electronic mapping platform.

**Figure 12.**
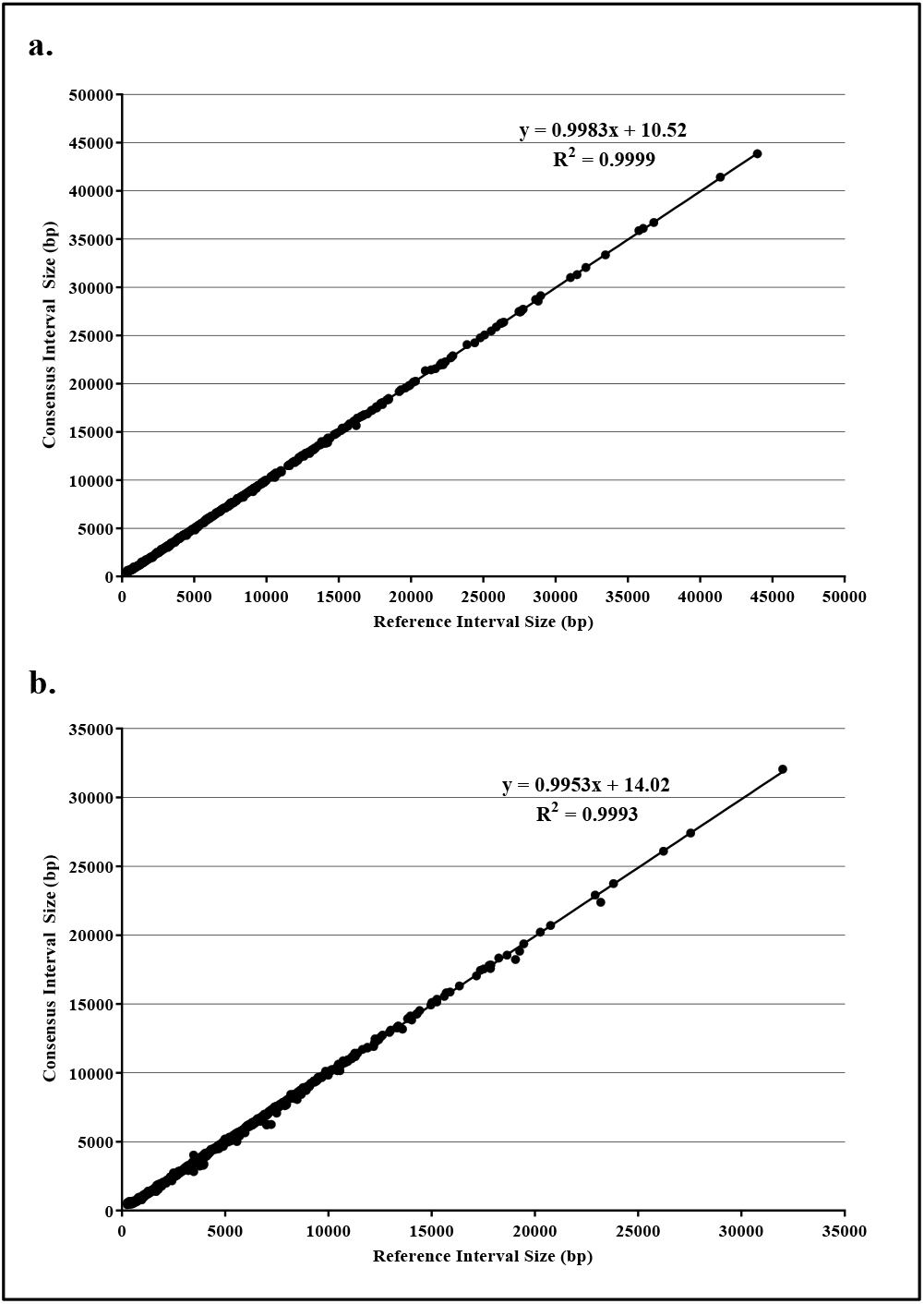
Plots of consensus vs. reference interval sizes for (a) Nt.BspQI and (b) Nt.Bpu10I mapping data sets with linear regression analysis indicated.

### *De novo* Assembly

The data from each nicking reaction can also be used for *de novo* assembly of maps. Assembly of the data from each nicking reaction was performed using proprietary assembly software. The assemblies covered 99.5% and 99.7% of the genome with an average depth of coverage of 250-fold and 180-fold for Nt.BspQI and Nt.Bpu10I respectively. The assemblies contained 2 and 8 contiguities respectively.

A screenshot of a 150 kb region of the assembly showing a portion of the reads used to create the assembly is shown in Figure 13. The software allows the user to view the entire genome map and expand any region of interest for more in depth observation. Statistical details of any interval in the assembly are available by using the cursor to click on the interval. The reads used to assemble the map are shown under the map and selecting an individual read reports the molecule length, false positive and false negative counts. The data screen also shows probe and interval coverage depth for the map.

**Figure 13.**
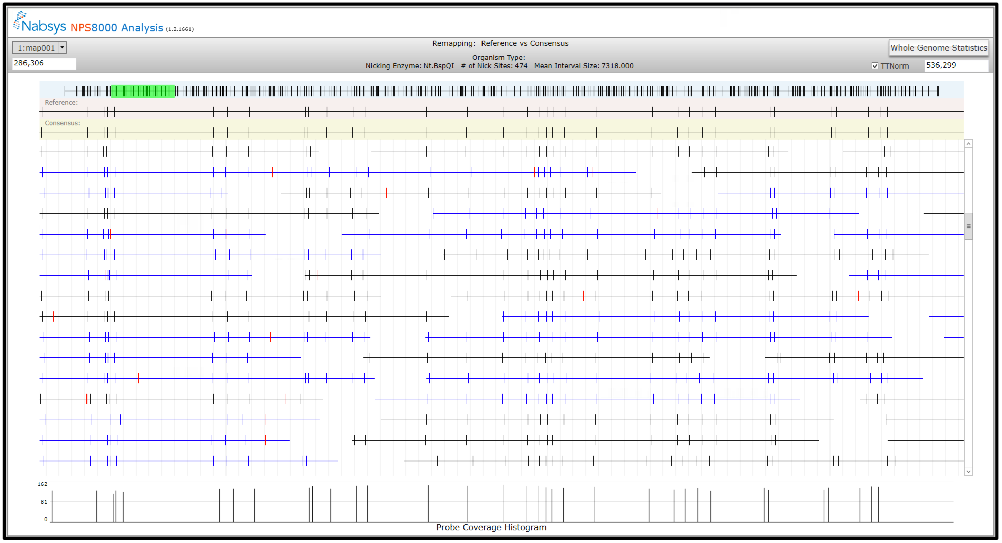
Screenshot of a portion of the reads used to assemble the Nt.BspQI map of *E. coli* MG1655. The assembled map is represented by the top line and the region highlighted in green is expanded immediately below the genome map. A subset of the reads used to assemble the highlighted region is shown. The entire set of reads may be observed by using the slider on the right side of the display. The bottom of the screen shows the coverage depth for each probe in this region of the assembly.

The reference genome of *E. coli* MG1655 was used to construct a reference map. An alignment of the Nt.BspQI assembly of *E. coli* MG1655 to the reference map is shown in Figure 14.

**Figure 14.**
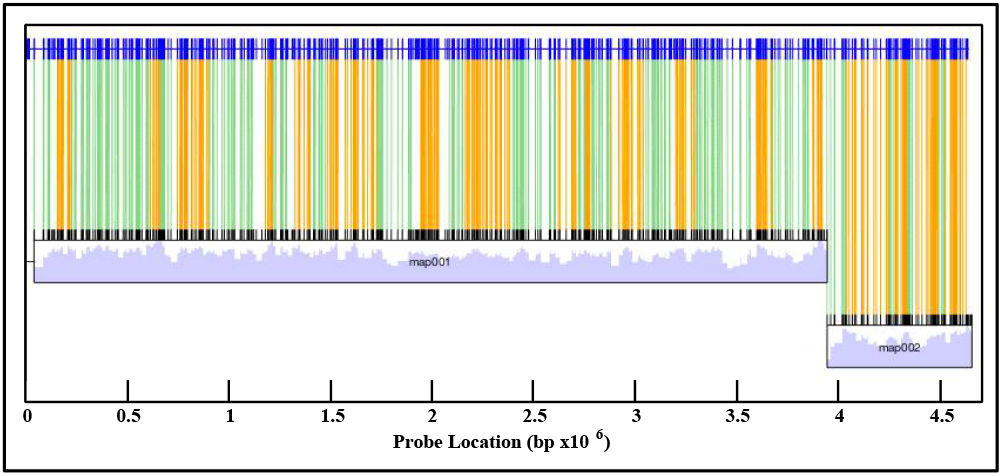
Alignment of the Nt.BspQI assembled map with the reference sequence. The reference map is displayed in dark blue and the assembled map is shown in black with probe alignments shown with green and orange lines. The depth of coverage of the assembly is shown in light blue below the assembled map.

The interval sizes in each assembled map were compared to the expected interval sizes in the reference map. The interval size comparisons for both the Nt.BspQI and Nt.Bpu10I assemblies are shown in Figure 15. Consistent with the remapping results, the assembly interval sizes are in close agreement with interval sizes determined from the reference sequences.

**Figure 15.**
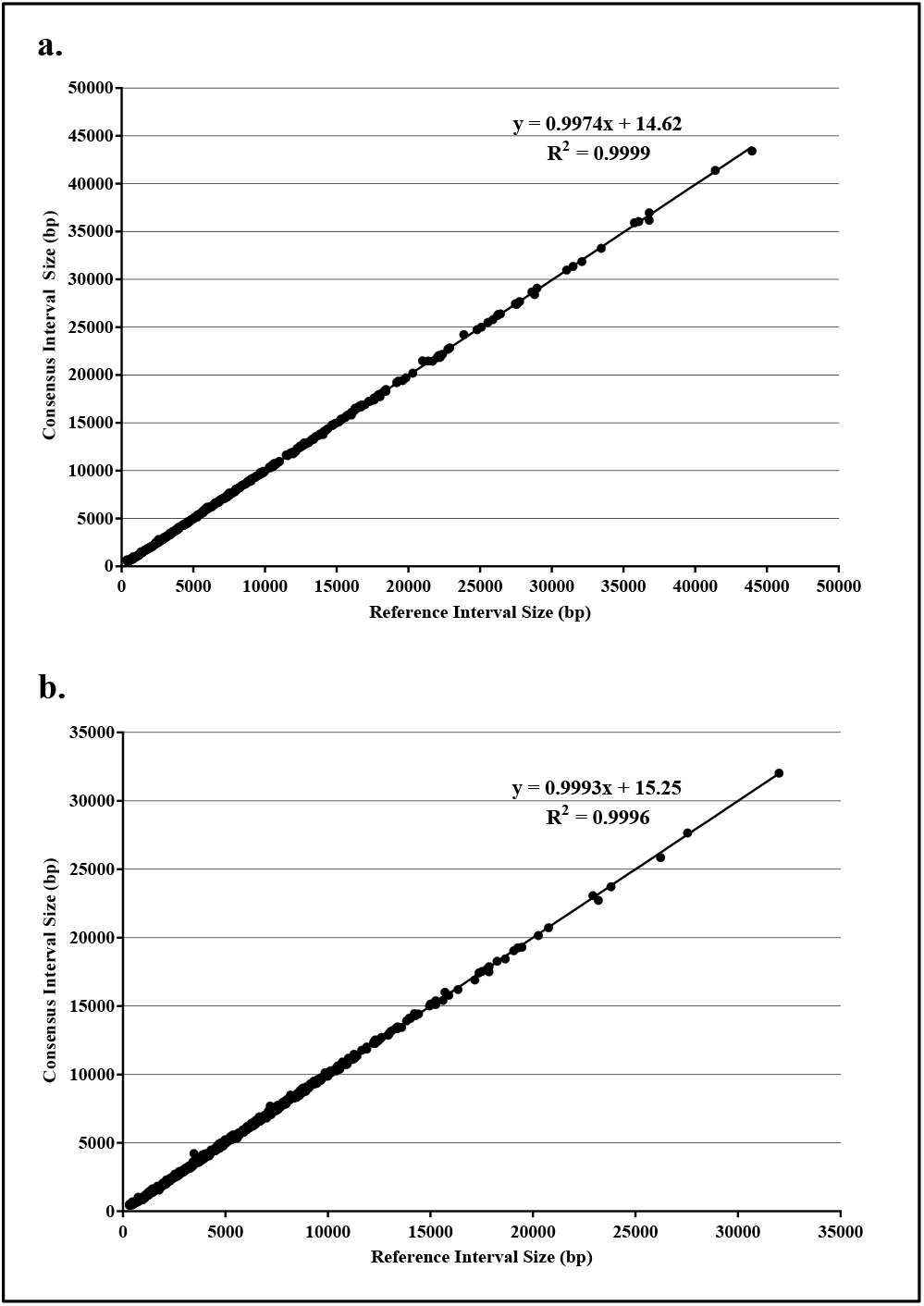
Plot of consensus vs. reference interval sizes for *de novo* assemblies. The assembly interval size vs. expected interval size is plotted for a) Nt.BspQI and b) Nt.Bpu10I maps of *E. coli* MG1655. A linear regression analysis is shown for the entire set of intervals.

## Conclusion

This report describes the design and performance of a unique and highly sensitive electronic detection scheme. The Nabsys platform is capable of detecting tags on single molecules of DNA and accurately measuring the distance between neighboring tags. These data can be used for applications relying on remapping ranging from next generation sequence contiguity validation to pathogen strain identification. The data can also be *de novo* assembled to produce dense, highly resolved maps suitable for scaffolding NGS contiguities and the detection of structural variants. The nature of the detection scheme obviates many of the physical limitations imposed by single-molecule optical detection and is capable of detecting interval sizes that are below the diffraction limit while molecules translocate through the detector at greater than 1 Mbp/s.

The higher resolution and precision of Nabsys whole genome mapping offers many advantages over existing technologies. Sample length requirements are relaxed with respect to optical methods because of the higher information content. In addition, higher density maps can be assembled from a wider variety of nicking enzymes than are available for optical mapping approaches.

Ongoing improvements will reduce the minimum size of intervals that are assembled by the platform, further increase the precision of interval determination, and increase platform throughput.

## Materials and Methods

### Reagents

Lambda DNA, Nt.BspQI Nt.Bpu10I, and 10X NEBuffer 3 were purchased from New England Biolabs (Ipswich, MA). RecA protein was purchased from Enzymatics Inc. (Beverly, MA). *E. coli* was purchased from ATCC (Manassas, VA). LB broth media was purchased from Teknova (Hollister, CA). NucleoBond AXG 20 columns and NucleoBond Buffer Set III were purchased from Macherey-Nagel, Inc. (Bethlehem, PA).

### DNA Isolation

A single *E. coli* MG1655 colony was selected for inoculation in 5 mL standard Luria Bertani (LB) medium. The culture was grown at 37°C with shaking at 250 rpm overnight until an OD_600_ of ~2.0 was achieved. *E. coli* genomic DNA was isolated using the Macherey-Nagel NucleoBond AXG 20 column system in conjunction with Macherey-Nagel NucleoBond Buffer Set III.

### DNA Tagging and Coating Reactions

The Nt.BspQI nicking reaction was performed by incubating purified *E. coli* DNA with Nt.BspQI at 4.4 U/*μ* g DNA in 1X NEBuffer 3. Reactions were carried out at 50°C for 1 h followed by 20 min at 80°C. The Nt.Bpu10I reaction was performed by incubating purified *E. coli* genomic DNA with Nt.Bpu10I at 2.7 U/*μ* g DNA in 1X NEBuffer 3. Reactions were carried out at 37°C for 1 h followed by 20 min at 80°C. Nabsys proprietary tags were attached by incubating the nicked DNA with the tag at room temperature for 30 min. The sample was then coated with RecA protein in the presence of ATPγS at 37°C for 2 h.^40^

